# Human Respiratory Syncytial Virus-induced immune signature of infection revealed by transcriptome analysis of clinical pediatric nasopharyngeal swab samples

**DOI:** 10.1101/2020.05.20.106492

**Authors:** Claire Nicolas De Lamballerie, Andrés Pizzorno, Julia Dubois, Blandine Padey, Thomas Julien, Aurélien Traversier, Julie Carbonneau, Elody Orcel, Bruno Lina, Marie-Eve Hamelin, Magali Roche, Julien Textoris, Guy Boivin, Catherine Legras-Lachuer, Olivier Terrier, Manuel Rosa-Calatrava

## Abstract

Human Respiratory Syncytial Virus (HRSV) constitutes one the main causes of respiratory infection in neonates and infants worldwide. Transcriptome analysis of clinical samples using high-throughput technologies remains an important tool to better understand virus-host complex interactions in the real-life setting but also to identify new diagnosis/prognosis markers or therapeutics targets. A major challenge when exploiting clinical samples such as nasal swabs, washes or bronchoalveolar lavages is the poor quantity and integrity of nucleic acids. In this study, we applied a tailored transcriptomics workflow to exploit nasal wash samples from children who tested positive for HRSV. Our analysis revealed a characteristic immune signature as a direct reflection of HRSV pathogenesis and highlighted putative biomarkers of interest.

## Background

Respiratory Syncytial Virus (HRSV) constitutes one of the main causes of respiratory tract infection in newborns and young children worldwide [1] but also in the elderly, immunocompromised, and patients with chronic heart and lung conditions [2]. The global HRSV disease burden is estimated at approximately 200,000 deaths and more than 3 million hospitalizations per year [1,3]. Despite numerous attempts and ongoing clinical trials, no efficacious HRSV vaccine is yet available, and the specific therapeutic arsenal currently available is very limited and remains relatively expensive [4]. In this context, we urgently need to increase our understanding of HRSV pathogenesis and the multiple facets of its virus/host interactions.

Much of the HRSV-induced disease is considered as the reflection of the host innate immune response to infection [5,6], with respiratory epithelial cells and monocytes/macrophages being the main actors in this response [7,8]. Indeed, HRSV infection was previously shown to up-regulate the expression of host genes involved in the antiviral and cell-mediated immune responses, such as genes coding for interferons (IFNs) and more largely several cytokines/chemokines such as CXCL10/IP-10, CXCL8/IL-8, MCP-1/CCL2, RANTES/CCL5 or IL6 [8]. For example, we previously demonstrated that HRSV infection, alone or in the context of bacterial co-infection, strongly promotes CXCL10/IP-10 expression in human macrophages [9]. We also showed that HRSV infection of human respiratory epithelial cells induces a strong disequilibrium in the p53/NF-kB balance, which appears to contribute to the up-regulation of several proinflammatory cytokines and chemokines [10]. One limitation of these *in vitro* approaches is that they do not necessarily reflect the whole complexity of the *in vivo* environment. In this context, we advantageously investigated the HRSV-induced host response using an innovative and highly relevant primary human reconstituted airway epithelial model, cultivated at the air-liquid interface to assess previously undescribed facets of the HRSV biology, such as the impact of the infection on cilium mobility and morphogenesis [11].

The development of high-throughput “omics” approaches has contributed to deepen our understanding of the multiple levels of interplay between respiratory viruses and the host cell [12–15]. These approaches, in addition to be very informative about the dynamic interplay between the virus and the host and hence the pathogenesis mechanisms, could also constitute a powerful tool to identify new therapeutic targets and/or propose novel antiviral strategies. In the case of HRSV, few studies have investigated the transcriptomic host response using clinical specimens, and an even more limited number have exploited respiratory tract samples [16–18]. A major challenge associated with transcriptome analysis of clinical samples is the intrinsic low copy number and/or low integrity of the nucleic acids recovered. To tackle these hassles, several research groups, including ours, have proposed and developed adapted/optimized sample processes [19–21].

In this study, we investigated the impact of the infection on the host cell using nasal washes from hospitalized children with lab-confirmed HRSV infection. Samples were processed with adapted protocols and transcriptomic signatures were obtained by hybridization on the HuGene 2.0 st Affymetrix microarray and subsequent process of the data. We compared our results to published pediatrics blood microarray datasets for the establishment of a nasal-specific signature. We also included biological results obtained using our previously described relevant human reconstituted airway epithelial (HAE) model of HRSV infection [11] for a deeper comprehension of the virus impact on the host epithelium. The analysis of HRSV-induced gene expression signature validated the importance of several IFN and cytokine-related pathways, in line with previous studies, but also provided valuable insight on potential biomarkers of diagnostic interest or as surrogates for the evaluation of future innovative treatments.

## Methods

### Clinical samples and ethical considerations

Written consent was obtained from parents of the three hospitalized children with lab-confirmed HRSV infections. Control samples come from the collection of samples established by the Québec CHU in the context of RespiVir surveillance study. The protocol was approved by Ethics committee of the CHU de Québec-Université Laval. Nasal wash samples were collected in RNAlater® Stabilization Solution (Thermo Fisher Scientific).

### RNA extraction and microarray experiment

Isolation of total RNA from nasal washes was performed using the RNeasy Micro kit (QIAGEN) with Dnase I treatment following the manufacturer’s instructions. Samples were quantified using the Quantifluor RNA System (Promega) and qualified using Agilent RNA 6000 Pico chip on Bioanalyzer 2100 (Agilent Technologies) according to manufacturer’s instructions. Whole RNA amplification using three rounds of in vitro transcription was then performed in two steps. First, the ExpressArt Trinucleotide mRNA amplification Pico kit (Amp Tec) was used for RNA amplification using trinucleotide and IVT transcription with a minimal input requested of 100pg. Then, ss-cDNA synthesis was performed with the GeneChip® WT PLUS Reagent Kit (Affymetrix) with a minimal input requested of 5.5 μg. cRNA and ss-cDNA quality control was assessed (Nanodrop and Bioanalyzer). Labeled cRNA was hybridized to GeneChip Human Gene 2.0 ST Array (Affymetrix) for 16h at 45°C and scanned using the Affymetrix 3000 7G Scanner..CEL file generation and basic quality controls were performed with the GeneChip™ Command Console® (Affymetrix).

### Data analysis

Data were analyzed using the R software and its xps (eXpression Profiling System) package (version 1.32.0) downloaded from www.bioconductor.org. The source files (CLF, PGF and transcript files) were downloaded from the Affymetrix website (http://www.affymetrix.com/site/mainPage.affx). Quality controls were performed to assess technical bias, RNA degradation levels and background noise. Preprocessing steps consisted in background correction, RMA normalization, probe summarization, and log2 transformation. A linear model was used to assess differential expression with the limma (Linear Models for Microarray Data) R/Bioconductor software package [22]. Genes were considered for subsequent analysis if they exhibited at least a 2-fold change in expression levels compared to the control samples coupled with p-values < 0.05. In order to further functionally characterize the patient transcriptomic signature, the web-based tool DAVID 6.8 was used to determine the enriched pathways [23]. Genes predicted by TargetScan 7.2 [24] to be targeted by the up- or down-regulated miRNAs with cumulative weighted score < −0.5 were used for functional enrichment analysis using the same web-based tool. To further comprehend the connexions between the modulated genes in our study, we chose to represent the interactome as a graph where nodes correspond with proteins and edges with pairwise interactions using the web-based tool STRING **11.0 [30]**(https://string-db.org), paired with Markov Clustering (MCL [26]) in order to extract relevant modules from such graphs.

### Pediatric mRNA datasets

We chose three datasets from HRSV-infected host transcriptomic studies publicly available on Gene Expression Omnibus database (GEO) and ArrayExpress. Two of them used peripheral pediatric blood samples (GSE69606 & E-MTAB-5195) and the third one focussed on PBMC gene expression responses to infection (GSE34205; n=51 HRSV-infected & n=10 controls). We extracted raw data corresponding severe disease samples from the series GSE69606 (n=8) and MTAB-5195 (n=18) and the recovery corresponding samples or healthy control samples (each n=8). Raw data were processed as previously described and differential analysis was performed according to the same thresholds (p-value < 0.05 and absolute fold-change > 2). We then compared the subsequent gene lists for tissue-specific gene expression assessment.

### Reconstituted human airway epithelia (HAE) and viruses

To counter for ultra-low nucleic acid quantities, the MucilAir® human airway epithelia (HAE) from Epithelix SARL was used for validation purposes. These HAE were maintained in air-liquid interphase with specific MucilAir® Culture Medium in Costar Transwell inserts (Corning) according to the manufacturer’s instructions. As previously described, apical poles were gently washed with warm PBS and then infected with a 150-μL dilution of HRSV-A (Long) virus in OptiMEM medium (Gibco, ThermoFisher Scientific) at a MOI of 1. Control HAE were mock-infected in the same conditions with MucilAir® Culture Medium with OptiMEM as the inoculum. After 6 days, total tissue lysates were harvested and total RNA was extracted as previously described [11,21]. The Human Respiratory syncytial virus (HRSV-A Long strain ATCC-VR26) was produced in LLCMK2 cells (ATCC CCL7) in EMEM supplemented with 2 mM L-glutamine (Sigma Aldrich), penicillin (100 U/mL), streptomycin (100 μg/mL) (Lonza), at 37 °C and 5% CO2.

### NanoString nCounter validation

The nCounter plateform (NanoString technologies) was used for mRNA detection of a 86 gene panel, according to manufacturer’s instructions [27]. This custom panel gathers immunity-related genes (cytokine production, proliferation T cells, interferon-gamma-mediated signaling pathway, among others). 300 ng of total RNA were hybridized to the probes at 67°C for 18 hours using a thermocycler (Biometra, Tprofesssional TRIO, Analytik Jena AG, Jena, Germany). After removal of excessive probes, samples were loaded into the nCounter Prep Station (NanoString Technologies) for purification and immobilization onto the internal surface of a sample cartridge for 2-3 hours. The sample cartridge was then transferred and imaged on the nCounter Digital Analyzer (NanoString Technologies) where color-codes were counted and tabulated for the 86 genes. Counts number were normalized by the geometric mean of HPRT1 (NM_000194.1), DECR1 (NM_001359.1), RPL19 (NM_000981.3), POLR2A (NM_000937.2) and TBP (NM_001172085.1) housekeeping genes count number, as well as the negative and positive control values using nSolver analysis software (version 4.0, NanoString technologies). Gene expression results are expressed in fold change induction compared to the mock-infected condition.

## Results

### Differential gene expression in HRSV-infected samples

In this study, we assessed nasal airway gene expression on pediatric nasal wash samples (3 infected and 5 controls). Given the low quality and low integrity of these sensitive samples (quality status available in **Supplementary table 1**), previously published adapted protocols were successfully used for their amplification and their subsequent hybridization on an Affymetrix GeneChip™ Human Gene 2.0ST [19]. An overview of the customized sample process and workflow is presented in **Figure 1A** and the subsequent hierarchical clustering of all analyzed samples is featured in **Figure 1B.** Despite the known heterogeneity of clinical samples, HRSV-infected and non-infected samples clustered appropriately to their corresponding experimental group.

**Figure 1.**
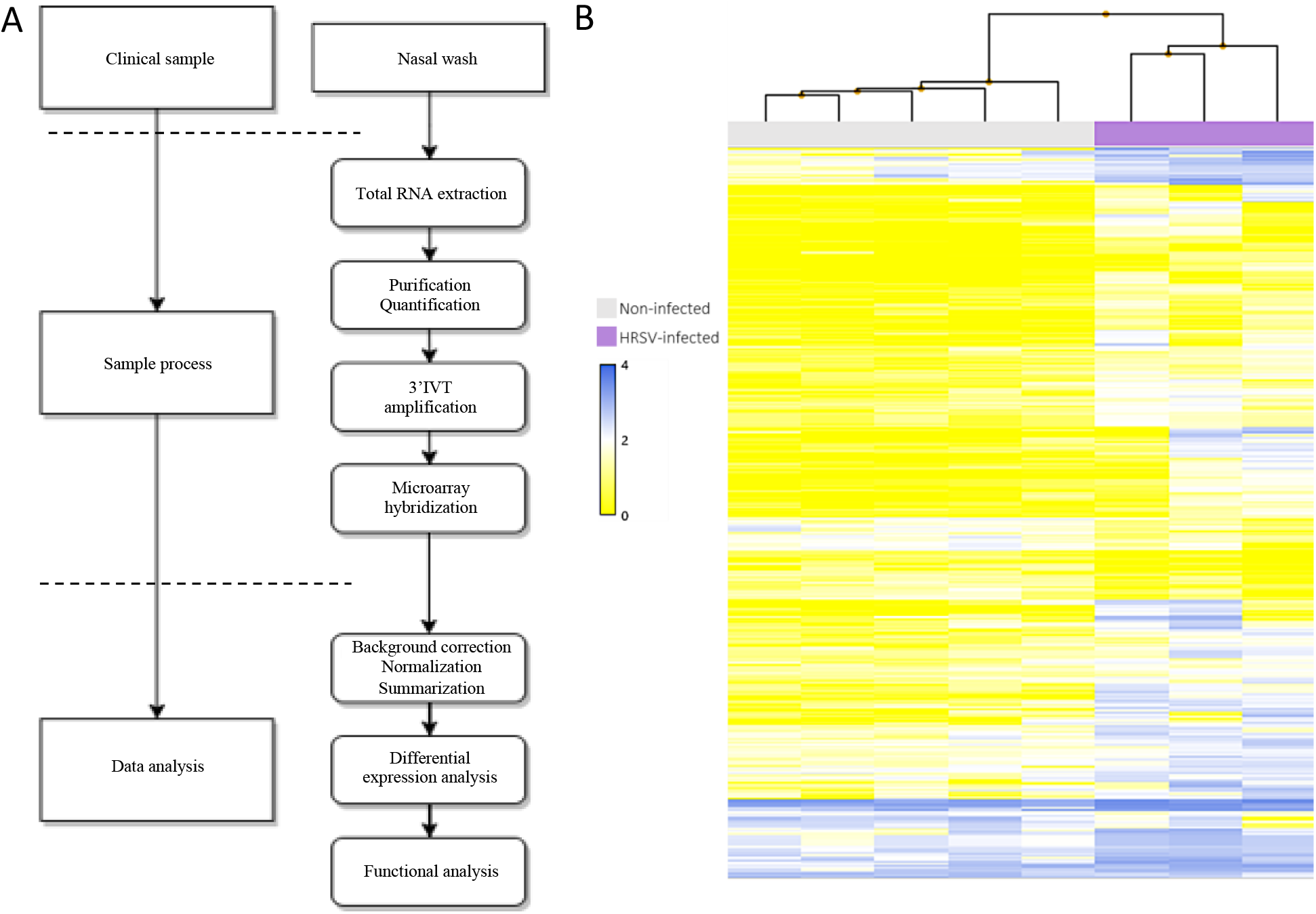
Sample processing workflow and transcriptomic hierarchical clustering. **(A)** Adapted workflow for the processing and exploitation of clinical samples with low RNA quality/quantity. **(B)** Hierarchical clustering of the signal intensities corresponding to all infected and non-infected clinical samples evaluated in the study. The resulting clusters are representative of the degree of similarity between samples and enable clustering into infected and non-infected experimental groups. The height of the y-axis at the branching points is a measure of similarity; y-axis units are arbitrary. For representation purposes, data was auto-scaled and log2 transformed.

For differential analysis, genes were considered significantly modulated if they exhibited at least a 2-fold change in expression levels compared to the control samples, with p-values inferior or equal to 0.05. Using these criteria, we listed a total of 296 differentially expressed genes, 258 (87.16%) of them being up-regulated (**Supplementary table 2**). This unbalanced up-*versus* down-regulated ratio was quite in line with previous observations [11]. As expected in the context of infected samples, many significantly up-regulated genes with fold changes far above 5 were related to the immune and IFN responses, such as ISG15, OASL, CXCL10/IP-10, CCL2-3, IFITM1-3 or IRF1 (**Supplementary table 2**). In contrast, among the 38 down-regulated genes, we listed genes associated with protein heterodimerization activity (SRGAP2C, HIST3H2BB and NTSR1), genes encoding zinc finger protein (ZNF439, ZNF28, ZNF286B, ZNF500) or transmembrane proteins acting as receptors or T-cell co-activators (such as SLC7A5P1, MSLNL or NTSR1). Of note, an important fraction (26%) of the down-regulated genes was represented by miRNAs, such as hsa-miR-572, hsa-miR-486-2, hsa-miR-1229 or hsa-miR-663b (**Supplementary table 2**). Aside from miRNAs, only 3 other down-regulated genes (KIR3DL2, SLC7A5P1 and SRGAP2C) had fold changes lower than −3.

### Gene Ontology-based functional enrichment analysis

To provide further functional interpretation of these clinical transcriptomic signatures, we then performed a Gene Ontology (GO)-based functional enrichment analysis using the web-based DAVID v6.8 toolkit (https://david.ncifcrf.gov/). GO terms, and particularly Biological Processes (BP), were considered enriched if their fold enrichment was higher than 2 and the Benjamini-Hochberg corrected enrichment p-value was inferior to 0.05. This BP enrichment was based on the global list of deregulated genes (**Figure 2**). As anticipated, the most enriched BP were primarily associated with interferon response (ex: GO:0060337; GO:0060333), response to virus (ex: GO:0009615; GO:0051607) or antigen processing (ex: GO:0002479; GO:0042612), which represent 16 out of the 24 most enriched BP listed (**Figure 2A**). Interestingly, the remaining GO terms were mainly related to mitochondria/respiratory burst (ex: GO:0005739; GO:0045730; GO:0045454) or ubiquitin ligase (ex: GO:0051437; GO:0051436).

**Figure 2.**
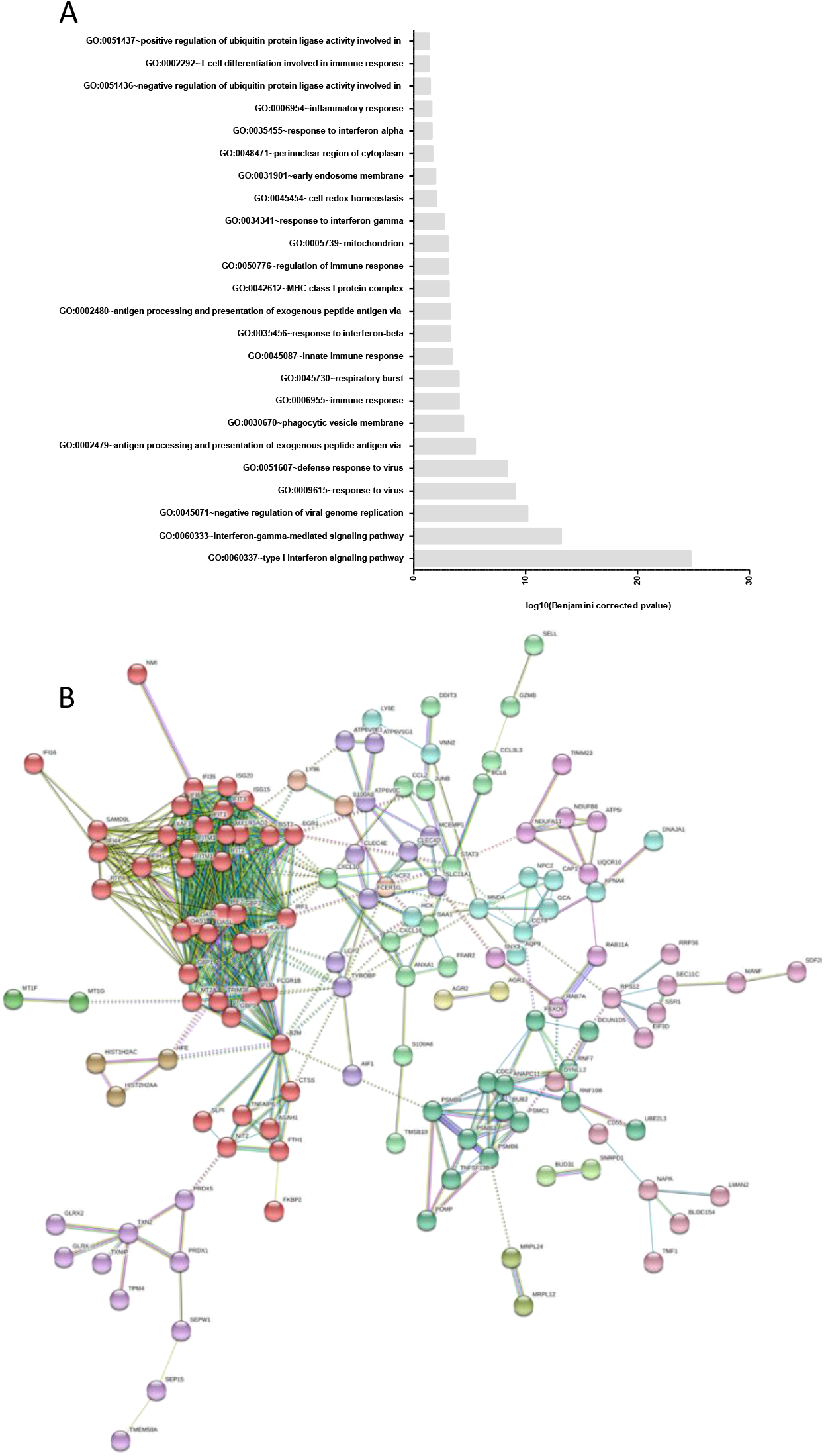
Gene Ontology-based functional enrichment and protein-protein interaction analyses. **(A)** Enriched biological process terms corresponding to the up-regulated gene list (enrichment score >2 and Benjamini-Hochberg corrected p-value <0.05). The down-regulated gene list did not present sufficient enrichment to pass our thresholds. **(B)** Evidence view of predicted protein associations associated with up-regulated genes in the HRSV-infected condition. Network nodes are host proteins and edges represent predicted functional associations. The color-coded lines correspond to the types of evidence supporting predicted associations (minimum required interaction score: 0.7). Node colors correspond to Markov clusters (MCL, inflation parameter: 1.5).

To better illustrate the impact of HRSV infection on the host immunity-related genes, we used the list of up-regulated genes and explored the functional association networks of their protein products using the STRING [25] database (https://string-db.org). As presented in **Figure 2B**, this analysis highlighted a functional network based on 241 distinct proteins (nodes) and 637 protein-protein associations (edges). These associations, which highlight proteins sharing functions but not necessarily physical interaction, are categorized into 15 relevant clusters, among which 10 contained more than 3 proteins (each color = 1 cluster by Markov Clustering [26] or MCL). Two major hubs concentrating a large number of edges were identified. The main hub consisted of proteins related to the immune response, with a central place for major actors like CXCL10, OASL and ISG15 (**Figure 2B**, red dots). The second hub harbored proteins involved in the positive and negative regulation of ubiquitin-protein ligase activity during the mitotic cell cycle (GO:0051436 and GO:0051437, medium purple dots). This includes any process that activates, maintains or increases the rate of ubiquitin ligase activity that contributes to the regulation of the mitotic cell cycle phase transition and vice versa. Much as the first hub would have been expected in an infectious context, the specific modulation of genes related to respiratory burst, cell redox homeostasis or ubiquitin-protein ligase activity by HRSV infection has been less studied.

Because of the numerous miRNAs deregulated, we used a prediction algorithm (TargetScan 7.2 [24]) for the identification of all genes targeted by the down- and up-regulated miRNAs. The predicted genes targeted by the down-regulated miRNAs were related to biological processes such as phagocytosis (GO:0006909), peptide cross-linking (GO:0018149) or positive regulation of release of cytochrome c from mitochondria (GO:0090200). Interestingly, the predicted targets of the up-regulated miRNAs are linked to similar processes: negative regulation of nucleic acid-templated transcription (GO:1903507) and negative regulation of T cell proliferation (GO:0042130) for the most enriched BPs (**Supplementary Figure 1**).

### Tissue-specific gene expression

In order to describe tissue-specific HRSV infection signatures, differentially expressed gene lists were extracted from three pediatric mRNA array datasets [28–30] and compared to our data to highlight similarities and differences between blood and respiratory airway transcriptional profiles, respectively (**Figure 3**). As previously shown, the overlap between the blood/PBMC and respiratory tract gene expression is scarce [29]. Only 6 genes (ANXA3, FCGR1B, OASL, BCL2A1, CLEC4D, RSAD2) were up-regulated in all analyzed datasets, mostly associated with the host immune response to the infection. As expected, both studies on peripheral blood shared a specific signature composed of 228 genes, whereas 51 additional genes were also modulated in PBMCs. These genes are mostly associated with the innate immune response of the host (GO:0045087). When comparing these signatures with the list of genes deregulated in our study, we highlighted 242 genes exclusively modulated in nasal washes, hence constituting specific drivers of the nasal epithelium signature. Among these tissue-specific modulated genes, genes were associated with the immune response regulation (GO:0050776) and, more precisely, with type I interferon signaling pathway (GO:0060337), or antigen processing and presentation (GO:0002479).

**Figure 3.**
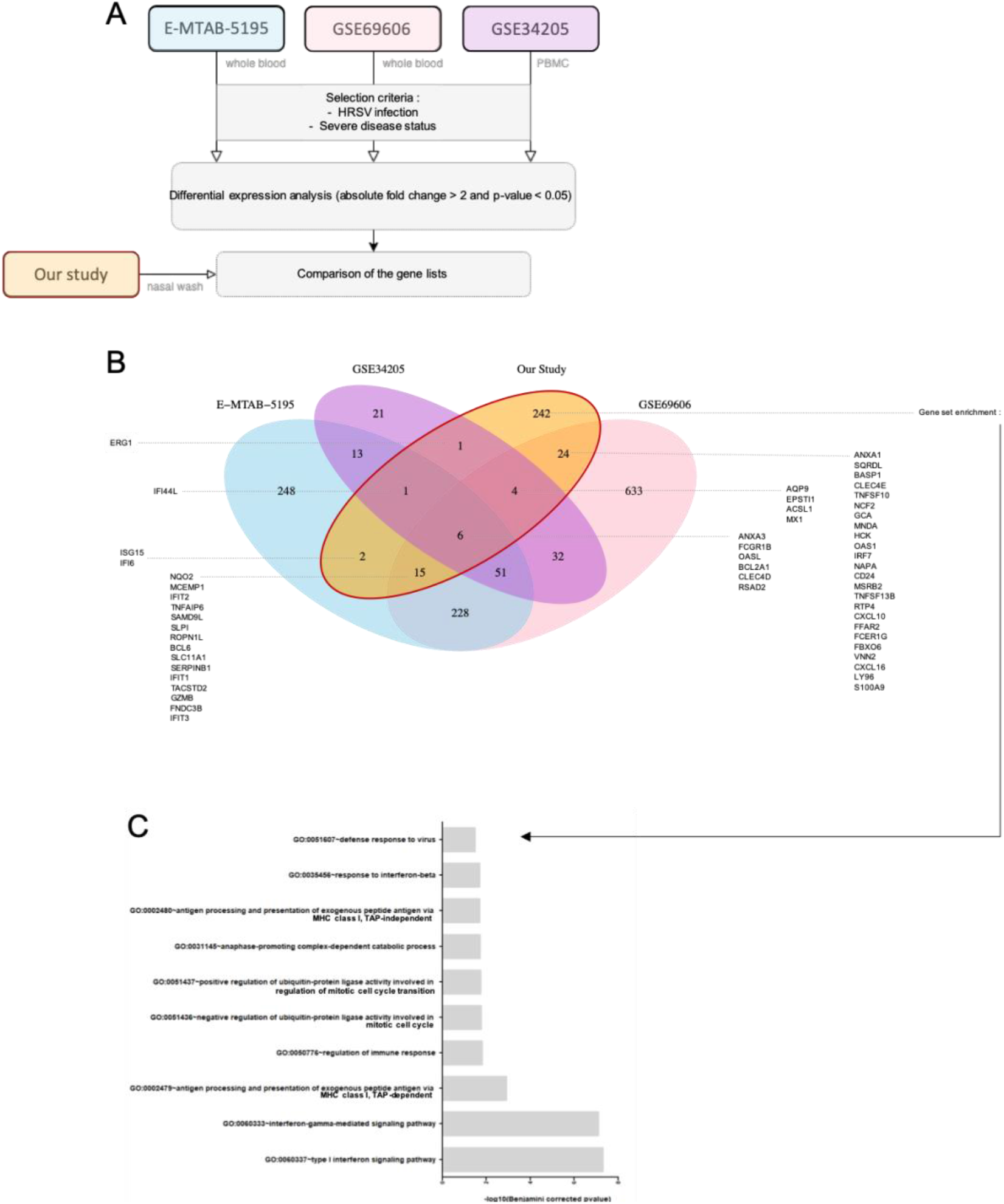
Gene expression cross-analysis as a key to tissue-specific local reaction to infection. **(A)** We selected 3 pediatric mRNA expression datasets for gene expression comparison across tissues. The GSE69606 dataset combines samples from 26 patients with acute HRSV infections, with symptoms spanning from mild to severe and the corresponding recovery paired samples. The E-MTAB-5195 dataset was originally used to investigate blood transcriptomics of 39 children during HRSV infection and for a longitudinal analysis to determine an 84-gene prognosis signature discriminating hospitalized infants with severe HRSV disease from infants with mild symptoms. The GSE34205 dataset is part of a wider study (GSE32140), aiming at establishing the signature induced by influenza and HRSV on PBMCs and primary airway epithelial cells. Because of the disease status of our study samples, we focused on severely ill children infected by HRSV. **(B)** Comparative cross-analysis of the gene lists on the 4 datasets (3 external plus ours). Common and specific infection features and gene enrichment analysis applied to the 242 nasal-specific genes are shown.

### Validation of differential expression in reconstituted human airway epithelium (HAE)

We then sought to validate these results in the context of experimental infections with the prototype HRSV A Long strain (MOI = 1) in a human reconstituted airway epithelial (HAE) model, as previously described [11]. This HAE model, issued from healthy donor biopsies, is composed of human primary ciliated columnar cells, mucus-secreting goblet and basal cells cultivated at the air-liquid interface, and has been successfully used to study viral infections and to evaluate the antiviral activity of many compounds in previous studies [11,21]. After 6 days of infection, HAE were lysed and total RNA was extracted and subsequently analyzed with the NanoString nCounter platform using a customized 94 immunity-related (cytokine production, T cell proliferation, interferon-gamma-mediated signaling pathway, etc) gene panel [27]. As shown in **Figure 4**, 39 out of the 94 genes in the NanoString panel were differentially modulated in the HRSV-infected condition compared to the mock-infected control. Despite the differential nature of infectious samples, the comparison of global gene expression modulation results between Affymetrix microarray (clinical samples) and NanoString (experimental infections in HAE) assays showed a correlation coefficient of 0.63. Unsurprisingly, the most up-regulated gene in the infectious context is CXCL10/IP-10, followed by IFI44L, IDO1 and TNFSF13B, with expression ratios above 50 (**Figure 4**). The top 20 modulated genes are strongly linked to “type I interferon signaling pathway” (GO:0060337) or more widely to “response to virus” (GO:0009615 or GO:0051607). Altogether, the gene expression results observed in clinical samples were cross-validated using an alternative method and underline a global deregulation of the biological defenses of the host, notably in the case of interferon stimulated genes (ISGs) that constitute a hallmark of many infectious and/or autoimmune disease states.

**Figure 4.**
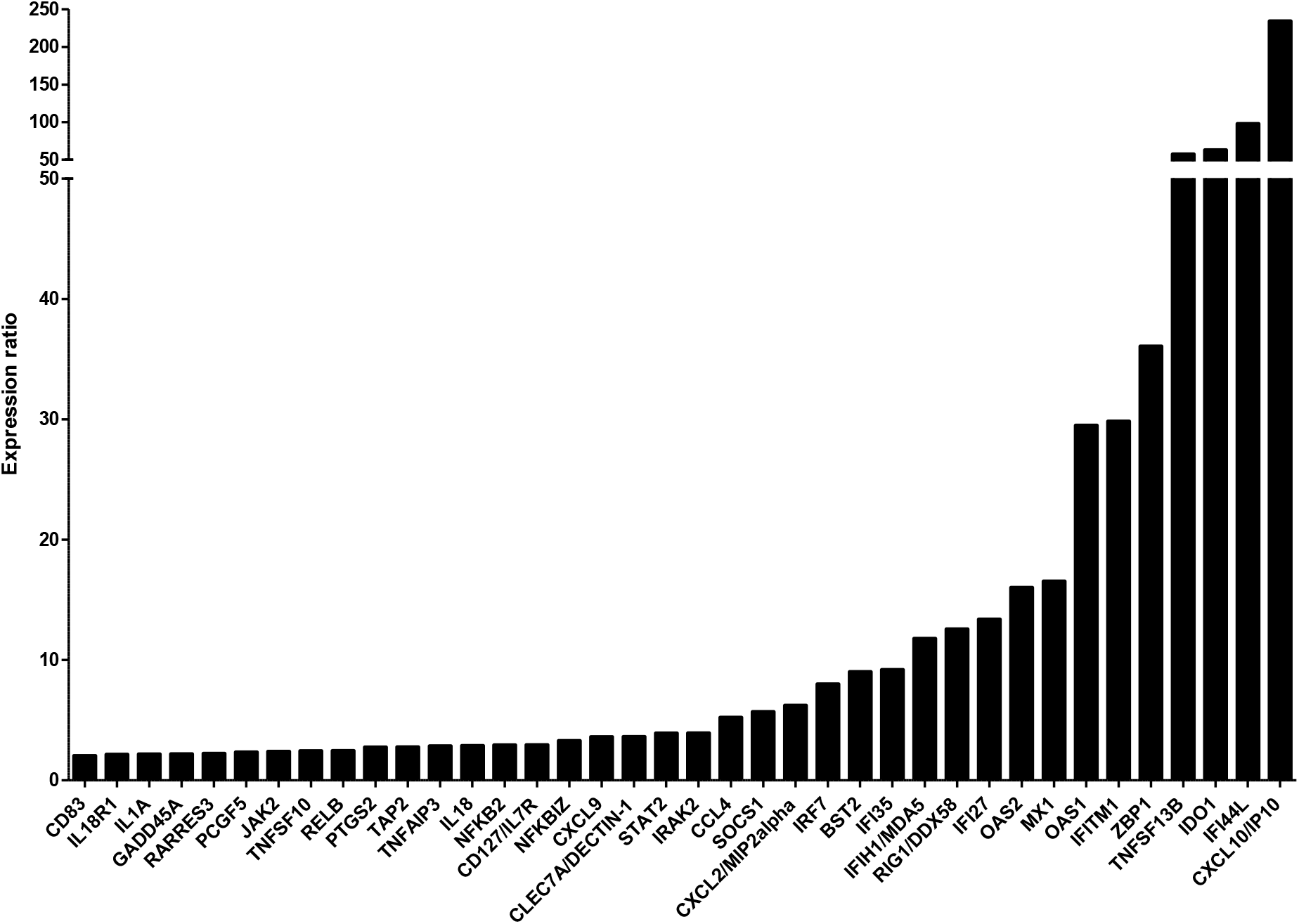
Experimental validation of gene expression results by NanoString assay in human airway epithelia (HAE). The expression of immunity-related genes in infected nasal HAE was validated using Nanostring nCounter technology. Data processing and normalization were performed with nSolver 4.0 analysis software and significant results (absolute fold change > 2) are expressed in fold change induction compared to the mock-infected condition.

## Discussion

In the particular context of HRSV infections, most of the respiratory samples collected for clinical studies are issued from pediatric patients, from whom only limited amounts of material are obtained. Moreover, given the fact that the main purpose of patient sampling is usually clinical analysis rather than research, the low quantity and quality of exploitable material is far from being optimal. In this study, we showed that these hurdles for the exploitation of highly degraded clinical samples could be mitigated by using adapted protocols and microarray, such as the Affymetrix GeneChip Human Gene 2.0 ST. Thus, despite starting from a limited number of children nasal washes presenting an acute HRSV infection, our adapted sample-processing pipeline enabled the determination and characterization of robust pediatric HRSV-induced nasal transcriptome signatures.

We initially identified well known markers of HRSV infection, namely the upheaval of the immune cascade [15–17], notably highlighting the strong overturning of CXCL10/IP-10 gene expression (coding for C-X-C motif chemokine 10, also known as interferon gamma-induced protein 10). We then advantageously used a biologically relevant reconstituted human airway epithelia (HAE) model to reproduce and validate these results by NanoString nCounter assay. IP-10 has already been described as the most abundant cytokine in bronchoalveolar lavages collected from HRSV-infected patients [31]. Moreover, our group already showed in a human monocyte-derived macrophage (MDM) model that the expression of this gene is highly impacted in the infection context [9]. We also identified up-regulated genes seemingly commonly modulated in different infection scenarios. For example, HRASLS2 is highly induced in RV infection [32], FFAR2 promotes internalization during IAV entry [33], and IFI6 has pleiotropic functions in HRSV, Dengue, or hepatitis C virus infections [34–36]. Conversely, our analysis revealed TMEM190 and MCEMP1 as potential specific biomarkers of HRSV infection. Although, TMEM190 expression was largely decreased in small airway epithelium by smoking [37] and MCEMP1 was proposed as a biomarker for stroke prognosis [38], a particular expression profile of these genes has not yet been described on other viral infections, for which they underscore further study as potential biomarkers. A similar rationale supports the study of TIMM23, strongly up-regulated in our study. A previous study conducted by Zaas et al. [15] identified a panel of 15 genes specifically modulated in HRSV-infected adults, of which FCGR1B, GBP1, RTP4, RSAD2, ISG15, IFIT2 were also significantly deregulated in our study. Despite the different nature of the biological samples used (children nasal washes *versus* adult blood), the high degree of concordance observed between our results and theirs supports a distinctive HRSV infection signature.

Using a differentiated, stratified and functional human airway epithelium model *ex vivoin vitro*, we validated a selected set of genes with the NanoString nCounter technology. Of note, none of the genes had a significant contradictory variation in both experiments. The only genes down-regulated in the pediatric samples with a fold change close to 1 and up-regulated with a fold change superior to 2 in the HAE model were IL1A, JAK2, PCGF5, SOCS1 and STAT2. This apparent discrepancy is coherent given the divergent experimental conditions (different gene expression technologies, collection timing, and/or nature of the sample considering the study model lacking immune cells). In this context, the validation of the up-regulation of 39 major genes of the immune response by the NanoString nCounter assay in the nasal epithelium model constitutes a relevant confirmation of the local immune disruption induced by HRSV infection in the nasal tissue. Hence, it consolidates the growing interest of such an accurate model for the study of viral infections, especially considering the abovementioned sample limitations in pediatric-oriented infections.

Besides those genes related to the immune response, our transcriptomics results also highlighted the modulation of genes and pathways related to a global mitochondrion cellular process disruption. This suggests that HRSV infection could unsettle less described biological processes related to the cAMP cascade, the redox complexes of mitochondrial respiratory chain, namely the respiratory burst. It consists of the production of high levels of reactive oxygen species (ROS) as a mean to discard internalized particles or pathogens following infection and phagocytosis. Although, this impact on the respiratory chain is a less explored aspect of HRSV pathogenesis in the respiratory tract, a study from Bataki et al. [39] investigated whether HRSV can directly signal to activate neutrophil cytotoxic function or not in the context of infant bronchiolitis. They assessed that when challenging neutrophils with diafiltrated HRSV, they could detect a lower activation of oxidative burst than in those challenged with unwashed virus or with virus free supernatant. Besides HRSV infections, the respiratory burst is known to be defective in influenza-infected neutrophils or during co-infections [40,41]. The disruption of such metabolic process could be a first clue regarding prognostic evolutions of children infected by HRSV.

In addition, the biological interpretation of the 38 down-regulated genes is not as straightforward. Indeed, no biological process or function was significantly enriched in our study, only modulations of individual gene were highlighted. Some markers, already described in the context of other viral infections, were found in the top down-regulated genes. For instance, Epstein–Barr virus (EBV) and human cytomegalovirus (HCMV) respectively express the viral proteins EBNA-3A and UL18, playing a key role in the antiviral host response to infection by preventing EBV/HCMV-infected cells from NK cell-mediated cytolysis [42,43]. This protection is known to be mediated partially by the inhibitory NK cell receptor KIR3DL2, whose gene expression is strongly inhibited in our HRSV infectious context. The SLC7A5P1 gene is also predicted to be linked to NK cells (GO:0032825~positive regulation of natural killer cell differentiation) whereas SRGAP2C had already be linked to HRSV bronchiolitis. Interestingly, we also observed the significant downregulation of several miRNAs. Among them, some are already described in literature such as miRNA-572, prognosis biomarker for renal cell carcinoma and sclerosis [44,45], or miRNA-769, included in a miRNA panel for discrimination between *Mycobacterium tuberculosis* infected and healthy individuals [46].

Regardless of the studied tissue, HRSV is consensually described as a major disruptor of the host immune response [47–49]. Here, comparing our signatures with 3 other ones extracted from pediatric whole blood transcriptomic analyses [28–30], we highlighted the common deregulation of 7 genes, independently of the tissue, and interestingly, 242 genes that seem to be specific to nasal epithelium HRSV-induced gene expression. This type of experiment had already been experimented in mice by Pennings et al., across three murine tissues (lung, bronchial lymph nodes and blood) [50]. They only found 53 genes regulated in common between the three tissues, notably GBP1, GBP2, GZMB, IFI44L, IFIT1, IFIT3, IFITM3, IRF7 and RTP4 genes, also significantly modulated in our study. By contrast, they described the GO/UniProt functional terms “acute phase”, “chemokine cytokine activity” and “antigen processing” to be characteristically attached to the lung signatures. This last term was specifically enriched in our study (GO:0002479: HLA-H, PSMB6, PSMB3, PSMC1, HLA-C, HLA-B, HLA-E, PSMB9, B2M), even if the global epithelium signature seems to evolve around the host immune response to infection.

Collectively, the transcriptomic analysis of nasal wash samples highlights the qualitative importance of such clinical samples, particularly in the context of their limited availability. The results obtained with a complementary approach such as the reconstituted HAE greatly contribute to bridge the knowledge gap in the understanding of the specific effects of HRSV on the host respiratory tissue and pave the way for several so far undescribed avenues of investigation.

---

**Supplementary Table 2.** Gene expression results (*available upon request*)

## Acknowledgements

The authors want to thank Sophie Assant for her help with the NanoString nCounter assay and Epithelix (Switzerland) for its help with MucilAir® human airway epithelia (HAE). This work was funded by grants from Région Auvergne Rhône-Alpes (CMIRA N° 14007029 and AccueilPro COOPERA N°15458 grants), and Canadian Institutes of Health Research (N° 229733 and 230187). Claire Nicolas de Lamballerie was funded by National Association for Research in Technology (ANRT). Guy Boivin is the holder of the Canada Research Chair on influenza and other respiratory viruses. Funding institutions had no participation in the design of the study, collection, analysis and interpretation of data, or in the writing of the manuscript.

## Author contributions

CNDL, AP, BL, GB, CLL, OT, MRC participated to conception and coordination of the study. CNDL, AP, BP, JC, EO, TJ, AT, BL, MEH, MR, JT, GB, CLL, OT, MRC carried out the experiments and analysis of the results. CNDL, JD, OT, AP, MRC designed the study and wrote the manuscript.

## Competing interests

The authors declare they have no competing interests.

---

**Suppl. Table 1:** Minimal quantities requested for microarray hybridization and homogeneity between samples are respected post amplification. The 8 samples (3 infected and 5 non-infected) were quantified and qualified after total RNA extraction (Quantifluor RNA System, Promega). The low quantity/quality observed after purification determined the need for subsequent amplification of the samples for hybridization. Samples underwent 3 rounds of unbiased *in vitro* amplification and sufficient cRNA was obtained to be used as cDNA template. Minimal quantities requested for hybridization on Affymetrix Human Genechip™ 2.0 ST Array were reached for all samples after three rounds of *in vitro* transcription.

| Infection Status | Sample ID | Sample type | RIN after RNA extraction | Initial concentration quantification (ng/μL) | RIN after purification | Concentration after purification (ng/ul) | cRNA quantification after amplification and purification (ug) | cDNA quantity (ng/ul) after conversion | cDNA quantification after conversion (ug) |
| --- | --- | --- | --- | --- | --- | --- | --- | --- | --- |
| Non-infected | 1 | Total RNA | 2.6 | 0.68 | 1 | 0.11 | 7.835 | 166.78 | 5.3 |
| Non-infected | 2 | Total RNA | 2.6 | 0.67 | 1 | 0.27 | 5.225 | 205.78 | 6.6 |
| Non-infected | 3 | Total RNA | 2.6 | 1.6 | 1 | 0.084 | 4.92 | 130.11 | 4.2 |
| Non-infected | 4 | Total RNA | 1.2 | 0.27 | Not Available | 0.012 | 10.87 | 250.85 | 7.0 |
| Non-infected | 5 | Total RNA | Not Available | 40 | 1 | 1 | 16.54 | 314.3 | 8.8 |
| HRSV-infected | 6 | Total RNA | 2.6 | 0.790 | Not Available | 0.04 | 21.52 | 280.24 | 7.85 |
| HRSV-infected | 7 | Total RNA | 2.6 | 0.880 | 1 | 0.023 | 18.49 | 369.48 | 10.3 |
| HRSV-infected | 8 | Total RNA | 2.6 | 1.100 | 1 | 0.077 | 5.975 | 198.28 | 6.3 |

**Supp Figure 1.**
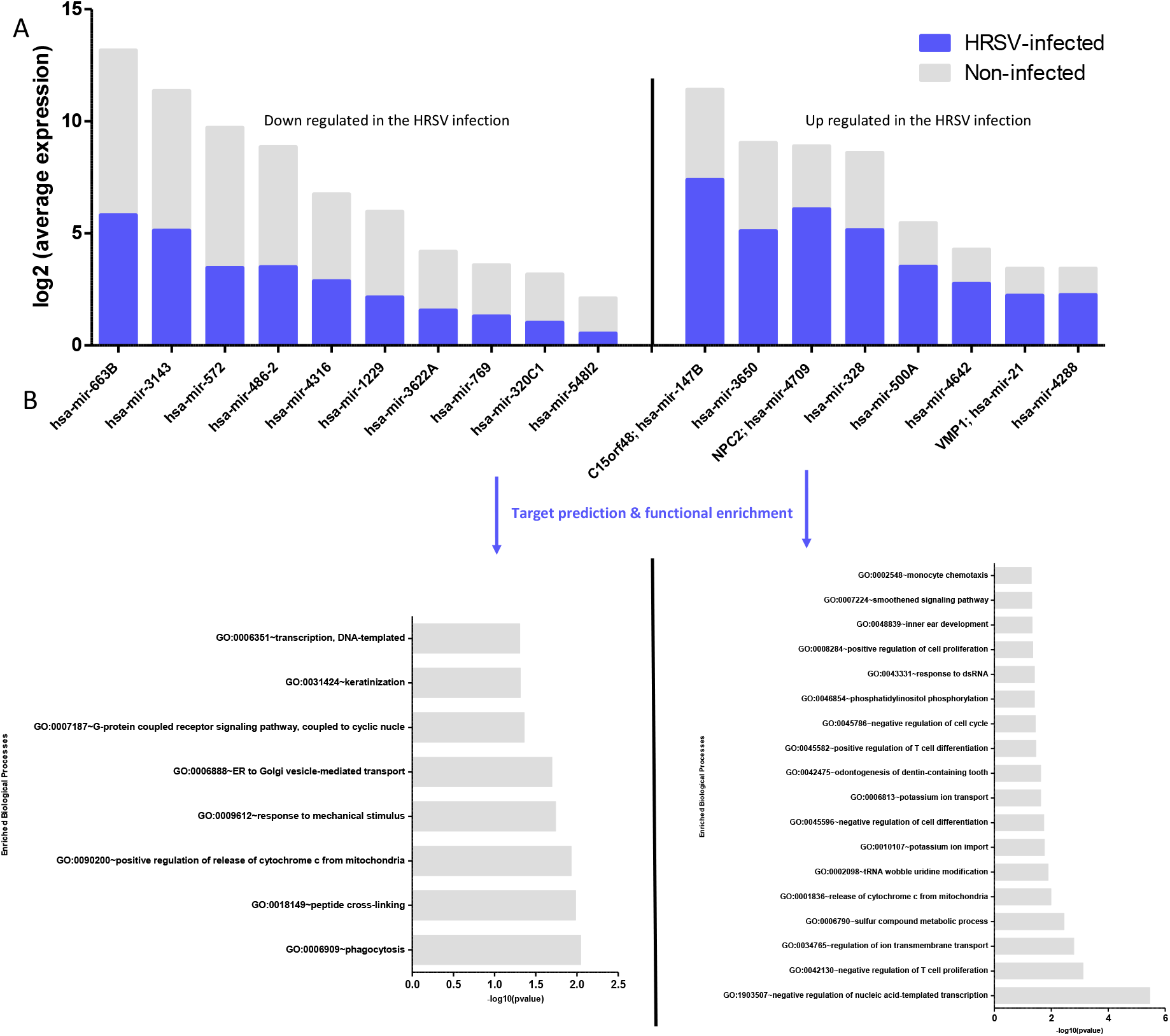
miRNA differential and functional analysis. **(A)** Log2 average expression values of both up- and down-regulated miR were extracted from the differential expression gene list (absolute fold change > 2 and p-value < 0.05). The identification of all genes targeted by at least one of the miRNAs was performed with the target prediction algorithm TargetScan 7.2. **(B)** Functional enrichment analysis (DAVID 6.8) of predicted biological targets to capture the involvement of such genes in several biological processes is shown

## Notes

### Competing Interest Statement

The authors have declared no competing interest.

